# SIREN: Suite for Intelligent RNAi Design and Evaluation of Nucleotide Sequences

**DOI:** 10.1101/2025.05.26.656188

**Authors:** Pablo Vargas-Mejía, Julio C. Vega-Arreguín

## Abstract

**Motivation:** RNA interference (RNAi) is a powerful tool for gene silencing across biological research, therapeutics, and agriculture. While siRNA design has benefited from advances in thermodynamic modeling and machine learning, comprehensive tools for designing long double-stranded RNAs (dsRNAs) with minimized off-target effects remain limited.

**Results:** Here, we present SIREN, an open-source Python pipeline designed to streamline RNAi construct design. SIREN integrates siRNA generation, thermodynamically-informed off-target prediction, scoring of dsRNA candidates based on cumulative off-target effects, and primer design for *in vitro* synthesis. It accepts user-defined transcriptomes for context-specific analysis and provides adjustable sensitivity settings balancing accuracy and computational demands. Benchmarking with plant, oomycete, and human transcriptomes demonstrates SIREN’s efficient scalability and the practical utility of medium sensitivity, recovering over 75% of high-sensitivity targets with significantly reduced computing times. Experimental validation in *Phytophthora capsici* confirms that SIREN effectively identifies highly specific RNAi constructs with no detectable off-target phenotypes in host plants.

**Availability and implementation:** SIREN is implemented in Python 3 and available under an open-source license at https://github.com/pablovargasmejia/SIREN; installer via PyPI: https://pypi.org/project/siren-rnai/.

## 1 Introduction

Since its discovery, RNA interference (RNAi) has become a crucial method for gene silencing, significantly advancing biological research, therapeutics, and agricultural pest control. RNAi primarily relies on double-stranded RNA (dsRNA) molecules processed into small interfering RNAs (siRNAs) that trigger the degradation of complementary mRNA, effectively silencing targeted genes.

Despite RNAi’s efficacy, designing dsRNAs and siRNAs involves significant challenges, particularly off-target effects where unintended genes may be inadvertently silenced, potentially causing undesirable outcomes. Although recent computational and machine-learning methods have enhanced siRNA design by integrating thermodynamic properties, nucleotide positional preferences (e.g., adenine enrichment), and chemical modifications (e.g., 2’-O-methylation), these advances have not been fully extended to comprehensive dsRNA design strategies (Monopoli *et al*., 2023; Neumeier and Meister, 2021). Current tools designed for dsRNA, such as si-Fi, rely on homology-based searches (Bowtie) and do not account for the cumulative off-target effects of multiple siRNAs derived from the same dsRNA (Lück *et al*., 2019). Tools like dsCheck consider cumulative effects but do not accept custom transcriptome inputs, restricting their utility to specific model organisms (Naito *et al*., 2005). Other available tools, such as dsRNAEngineer and dsRIP, also lack thermodynamic modeling and do not adequately account for cumulative off-target interactions (Cedden *et al*., 2025; Chen *et al*., 2025).

To address these limitations, we developed SIREN (Suite for Intelligent RNAi Design and Evaluation of Nucleotide Sequences), a bioinformatics pipeline especially designed to bridge this gap. SIREN systematically generates, evaluates, and ranks RNAi sequences, integrating thermodynamic models and comprehensive cumulative off-target assessment, providing researchers with a powerful tool to enhance precision in gene silencing applications.

## 2 Methods

### 2.1 Overview of the SIREN Pipeline

SIREN is a user-friendly Python pipeline that integrates multiple computational steps to facilitate efficient RNAi construct design and evaluation (general overview provided in Figure 1a-b).

**Figure 1.**
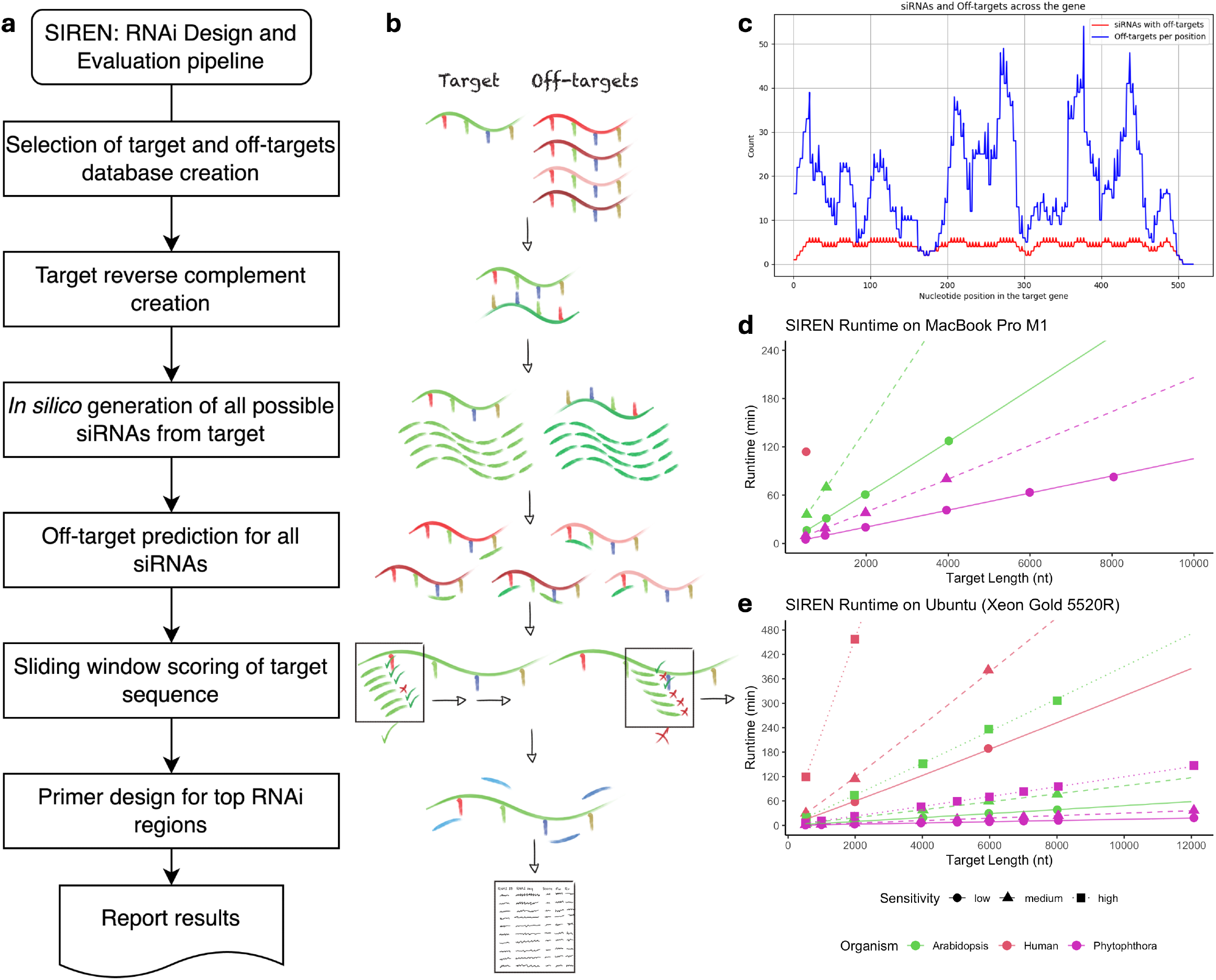
a SIREN pipeline overview. b SIREN schematic pipeline. c SIREN graphical output. d Runtime on a M1 MacBook Pro (8 cores used). e SIREN runtime over various databases on a Xeon Gold 5520R Ubuntu computer (46 cores used).

### 2.2 Input Processing and *in silico* siRNA Generation

SIREN initiates by accepting user-provided nucleotide sequences in FASTA format, which may include cDNA, mRNA, or CDS sequences. Alongside these sequences, the user specifies a target gene intended for silencing. First, SIREN generates a reference database of potential off-target sequences derived from the provided input, excluding the target gene to avoid biased predictions. The pipeline then computes the reverse complement of the target gene sequence to comprehensively identify all possible candidate siRNAs from both strands.

Subsequently, candidate siRNAs are generated systematically using a sliding window method. The density of generated siRNAs depends on the user-selected sensitivity option (high, medium, or low). Under high sensitivity, SIREN generates overlapping siRNAs at single-nucleotide increments along both strands of the target sequence. Medium sensitivity reduces computational load by generating siRNAs at increments of every four nucleotides, whereas low sensitivity (default setting) generates siRNAs every eight nucleotides. All generated siRNA sequences are stored in FASTA format (results_folder/other_files/sirnas.fa) with identifiers clearly indicating their positions of origin, such as sirna_1-21 or sirna_57-77_r for reverse-complementary sequences.

### 2.3 Off-target Evaluation

Default parameters (−e -25 -v 0 -u 0 -f 2,7 -p 0.01 -d 0.5,0.1 -m 60000) are selected to reliably identify biologically relevant off-target interactions. To optimize computational efficiency, SIREN utilizes parallel processing, splitting the off-target database into subsets distributed evenly across available CPU threads. The resulting RNAhybrid output is compiled into a single comprehensive file (results_folder/other_files/off_targets_results.txt), from which SIREN produces a summarized three-column table (results_folder/off_targets_summary.tsv). This summary ranks off-target transcripts according to the frequency of siRNA matches, listing each off-target gene, the number of matching siRNAs, and corresponding siRNA identifiers.

### 2.3 Off-target Visualization

The pipeline provides a graphical representation (Figure 1c) depicting the positional distribution of siRNAs associated with off-target matches along the target gene. This visualization effectively communicates regions of the target gene generating siRNAs with significant off-target potential.

### 2.4 Scoring, Selection and Primer Design

Following off-target assessment, SIREN compiles all possible RNAi sequences of user-defined length (default: 300 nt), constructed using four-nucleotide increments along the target sequence. Each RNAi candidate sequence is evaluated through a scoring algorithm considering two distinct penalties:

1. Off-target siRNA Penalty (−0.1 points): A penalty of −0.1 points is applied for each siRNA that fully matches an off-target transcript and is entirely contained within the evaluated RNAi sequence. Each siRNA contributes this penalty only once per RNAi sequence, regardless of how many off-targets it matches, preventing penalty inflation due to multiple off-target hits.
2. Redundant Off-target Hit Penalty (−30 per additional siRNA): If multiple distinct siRNAs within a single RNAi candidate target the same off-target transcript, an additional penalty of −30 points is applied for each extra siRNA beyond the first. This ensures strong penalization of RNAi sequences likely to cause extensive unintended silencing of a specific off-target gene.

Scores are calculated efficiently using parallelized computation across multiple CPU threads. From these results, the top 25 candidate RNAi sequences are reported, alongside shorter and longer variants (±50 nt), providing flexible options for experimental implementation. Optimal RNAi candidates undergo primer design via Primer3-py, with stringent considerations for melting temperature, specificity, and amplification efficiency. Final results, including RNAi sequences, associated scores, and primer pairs, are provided in the file results_folder/rna_sequences_with_scores.tsv (Figure 1a-b).

### 2.5 Benchmark and Testing

To assess SIREN’s computational performance, rigorous benchmarking was conducted across representative transcriptomes: *Arabidopsis thaliana* (TAIR10 from arabidopsis.org), *Phytophthora capsici* (LT1534 v11.0 from MycoCosm), and *Homo sapiens* (GRCh38 from ENSEMBL). The pipeline was tested on two distinct computational platforms: a MacBook Pro M1 with an 8-core ARM processor (MacOS Sequoia 15.4.1), and an Intel Xeon Gold 5220R workstation with 48 cores (Ubuntu 24.04.2 LTS).

From each reference database, twenty representative sequences ranging in length from approximately 500 to 12,000 nucleotides (0.5 kb, 2 kb, 4 kb, 6 kb, 8 kb, 10 kb, and 12 kb) were analyzed to evaluate computational efficiency and runtime scaling. Benchmarks were performed at all three sensitivity settings on the Xeon workstation, while only low and medium sensitivity levels were evaluated on the MacBook M1 due to hardware limitations. These benchmark analyses provided critical insights into runtime behavior, scalability, and practical performance considerations across varied computational environments and transcriptomic complexities.

### 2.6 Implementation and Availability

SIREN is developed in Python 3, integrating widely adopted open-source libraries: Biopython for sequence manipulation, RNAhybrid for off-target prediction, Primer3-py for primer design, and matplotlib for visualization. Complete source code, detailed documentation, and installation instructions are accessible on GitHub (https://github.com/pablovargasmejia/SIREN) and through PyPI (https://pypi.org/project/siren-rnai/), facilitating broad usability and reproducibility.

## 3 Results and Discussion

### 2.1 Performance

SIREN was benchmarked on two computational platforms: a MacBook Pro M1 (8-core ARM) and an Intel Xeon Gold 5520R workstation (46 cores used). Three transcriptome databases of increasing complexity were tested: *Phytophthora capsici* (20.4 Mb), *Arabidopsis thaliana* (64.9 Mb), and *Homo sapiens* (395.1 Mb).

On the MacBook M1, we ran only low- and medium-sensitivity modes due to hardware limitations. For *Phytophthora*, runtimes averaged ∼37 minutes for both low and medium sensitivity. For Arabidopsis, runtimes reached ∼59 minutes (low) and ∼53 minutes (medium), while a single *Homo sapiens* transcript required nearly 114 minutes at low sensitivity. When normalized by transcript length, the average processing time ranged from ∼0.61 seconds per nucleotide in *Phytophthora* (low) to ∼4.02 s/nt in Arabidopsis (medium), and peaked at ∼12.85 s/nt for the human transcript. These durations reflect not only the growing size of the reference databases but also the increased number of siRNAs generated from longer transcripts, each of which must be individually screened against the full target set. Despite the M1’s efficiency in general computing, its 8-core ceiling and lack of thread-level parallelism limit SIREN’s scalability under heavier workloads.

On the Xeon workstation, we evaluated all three sensitivity modes across all organisms. For Phytophthora, runtimes ranged from ∼3.4 minutes (low) to ∼7.2 minutes (high); for Arabidopsis, from ∼8.5 to ∼17.3 minutes. Human transcript processing ranged from ∼19.4 minutes (low) to ∼52.3 minutes (high), depending on transcript length and sensitivity mode. When normalized, runtimes on the Xeon averaged ∼0.15–0.33 s/nt, highlighting the substantial performance gain from parallel processing. Here again, longer transcripts took proportionally longer due to the increase in candidate siRNAs, but the server’s 46-core capacity allowed near-linear scaling even at high sensitivity (Figure 1d-e).

Crucially, more than 94 % of SIREN’s total runtime was consistently spent on the off-target search step, which relies on RNAhybrid—a widely used tool for miRNA and siRNA target prediction. RNAhybrid implements a thermodynamically grounded model of RNA-RNA interaction using dynamic programming, offering high biological accuracy but at the cost of speed. Because SIREN evaluates every candidate siRNA against every transcript in the reference set, runtime scales sharply with both the number of siRNAs and the size of the target database. This explains the particularly high runtimes seen for longer human transcripts under high sensitivity, which uses dense siRNA tiling.

Future versions of SIREN will prioritize acceleration of the off-target analysis phase, potentially through RNAhybrid batching, algorithmic optimization, or the use of approximate prefiltering methods. While alternative search strategies may offer speed improvements, RNAhybrid remains one of the most biologically reliable options for siRNA off-target prediction, and its continued integration ensures that SIREN maintains its accuracy and relevance for high-confidence RNAi design (Chen *et al*., 2019; Riolo *et al*., 2021).

### 2.2 Sensitivity levels

The three sensitivity settings differ primarily by the number of generated siRNAs: high sensitivity generates a maximum number ((target_length - siRNA_length)*2), medium sensitivity reduces siRNA numbers by approximately 75% (generating one every 4 nucleotides), and low sensitivity generates about 87.5% fewer siRNAs (one every 8 nucleotides). In this benchmarking exercise, we evaluated SIREN on twenty transcripts drawn from three distinct reference sets: Mycososm cDNA assembly of *Phytophthora capsici*, TAIR10 cDNA of *Arabidopsis thaliana*, and well-annotated human cDNA from Ensembl. For each target, the program was tested with the same conditions under each sensitivity setting (low, medium, high). When using low sensitivity, exact recovery of the 45 high-sensitivity designed RNAi ranged from 0 % to 67 % (median ≈ 20.6 %), whereas medium sensitivity recovered between 4 % and 91 % of those candidate sequences (median ≈ 48.9 %). In off-target screening, low sensitivity reported 232–9697 transcripts (median ≈ 2 517) versus 653–18175 (median ≈ 6 296) at high sensitivity; medium sensitivity called 426–16434 off-targets (median ≈ 4 463), consistently capturing a larger fraction of the high-sensitivity landscape. Allowing a ±10 % positional tolerance around each RNAi predicted sequence (sequences very likely falling within the same region), low sensitivity covered only 0 %–57.8 % of high-sensitivity target regions (median ≈ 28.9 %), while medium sensitivity reached 35.6 %–100 % coverage (median ≈ 76.7 %).

Regarding the most redundant off-target transcripts, those predicted to be bound by the largest number of siRNAs, we defined the top 20 % redundant off-targets as the subset of off-target transcripts whose “siRNA number” ranks in the upper quintile. This metric focuses on the highest-risk off-targets, since transcripts hit by multiple siRNAs are more likely to yield phenotypic noise. Under low sensitivity, only 12.8 % to 31.5 % (median ≈ 25.4 %) of these high-risk off-targets were recovered when compared to the high-sensitivity standard, indicating that low-sensitivity scans systematically underrepresent the most frequently targeted transcripts. Medium sensitivity substantially improved recovery (25.7 % to 51.5 %, median ≈ 48.7 %), nearly doubling the capture rate of critical off-targets and yielding Spearman ρ values up to 0.73 for redundant off-target counts.

Together with the exact-match and positional-tolerance assessments, the top-20 % redundancy analysis shows that medium sensitivity strikes a strong balance: it recovers a majority of the most problematic off-targets while requiring far fewer computational resources than a full high sensitivity run. In practice, medium sensitivity not only retrieves more of the high-priority off-target landscape but also stabilizes rank order among those off-targets that pose the greatest risk of confounding downstream assays.

### 2.3 Real RNAi Tests

SIREN was experimentally validated by silencing multiple *P. capsici* genes through application of dsRNA onto pathogen and host tissues (leaves and stems). dsRNAs were designed using SIREN against a combined transcriptome database of the phytopathogen *P. capsici* and four different plant hosts: tomato, chili pepper, cucumber, and melon. Following synthesis (cDNA amplification, T7 promoter fusion PCR, and T7-based in vitro transcription), none of the designed dsRNAs caused detectable off-target phenotypes in host plants, confirming practical effectiveness (Vargas-Mejía et al., manuscript in preparation).

## 4 Conclusions

SIREN efficiently facilitates the design and selection of highly specific RNAi constructs, balancing computational performance with sensitivity to off-target effects. Benchmarking demonstrates effective scalability, with medium sensitivity providing an optimal compromise between computational efficiency and accuracy. Experimental validation underscores SIREN’s practical utility, ensuring reliable RNAi design with minimal unintended off-target outcomes.

## Data availability

Source code is available at https://github.com/pablovargasmejia/SIREN, and installer is available at https://pypi.org/project/siren-rnai/

## Author contributions

Pablo Vargas-Mejía (Conceptualization, Software, Methodology, Writing-original draft, Writing-review and editing, Visualization), Julio C. Vega-Arreguín (Supervision, Writing-review and editing).

## Acknowledgments

This work forms part of the productivity of P.V.M. to his Doctoral degree from the Posgrado en Ciencias Biológicas program at Universidad Nacional Autónoma de México (UNAM). P.V.M. receives a doctoral fellowship from Secretaría de Ciencia, Humanidades, Tecnología e Innovación (SECIHTI). This study was financed with P.V.M. doctoral fellowship and by PAPIIT-DGAPA (project IN218024) to J.V-A.

